# Improving glycine utilization in *Escherichia coli*

**DOI:** 10.1101/2022.10.10.511530

**Authors:** Vincent Fung Kin Yuen, Daniel Zhi Jun Tan, Kang Zhou

## Abstract

*Escherichia coli* is a bacterium that has been widely used as host in industrial fermentation processes. Sugars and glycerol are currently used as feedstocks in most of such applications. To reduce the associated carbon footprint, there are many ongoing efforts in engineering the bacterium to utilize formate, a molecule that can be obtained from CO_2_ easily. Glycine is a key intermediate in a formate utilization pathway that has been reconstituted in *E. coli*. This study focuses on engineering *E. coli* to assimilate glycine into the central metabolism. We systematically compared three glycine utilization pathways and found that the glycine dehydrogenase pathway yielded the most stable strain. Through rational promoter engineering and evolution in a continuous stirred tank reactor (CSTR) with a mutator plasmid, we isolated a strain that was able to use glycine as the sole carbon and nitrogen source. It consumed 8 g/L glycine within 48 h. Whole genome sequencing revealed 40 changes in its genome, including a few in critical genes such as those encoding glutamate synthase and ATP synthase. The expression of the genes around the glyoxylate node was also found by RNA sequencing to be fine-tuned, presumably for reducing accumulation of the toxic aldehyde intermediate (glyoxylate). The strain obtained in this study could be useful in improving formate utilization in *E. coli*. The methods and equipment developed in this study (e.g., the customized, low-cost CSTR) could also facilitate training *E. coli* to utilize other non-conventional substrates.

## 1. Introduction

Glycine is a simple amino acid containing only two carbon atoms per molecule. It is an important intermediate in the reductive glycine pathway, which can be used to assimilate CO_2_ and formate into central metabolism of bacteria (Claassens et al., 2020; Yu & Liao, 2018). This pathway has been used in recent studies for enabling growth of *Escherichia coli* on CO_2_ and formate, which lays a solid foundation for developing carbon-neutral/negative biomanufacturing processes (Maia et al., 2021; Neuendorf et al., 2021; Van Peteghem et al., 2022). Despite of the central role of glycine in this pathway, to our knowledge, there is no study dedicated to improving *E. coli* growth on glycine.

Model wildtype *E. coli* strains are unable to use glycine as the sole/major carbon source. Three pathways have been explored for assimilating glycine into the central metabolism (Claassens et al., 2020; Yu & Liao, 2018). The first pathway condenses two glycine into one L-serine (**Figure 1a**), which is then deaminated by serine deaminase or threonine dehydratase to form pyruvate (Burman et al., 2004; Newman & Walker, 1982; Su & Newman, 1991; Su et al., 1989; Umbarger & Brown, 1957). The second and third pathways convert glycine into glyoxylate through glycine:alanine transamination or an oxidative reaction catalysed by a D-amino acid dehydrogenase (Claassens et al., 2020; Takada & Noguchi, 1985; Yu & Liao, 2018). Glyoxylate could be transformed into 2-phosphoglycerate through the glycerate pathway (**Figure 1b**).

**Figure 1.**
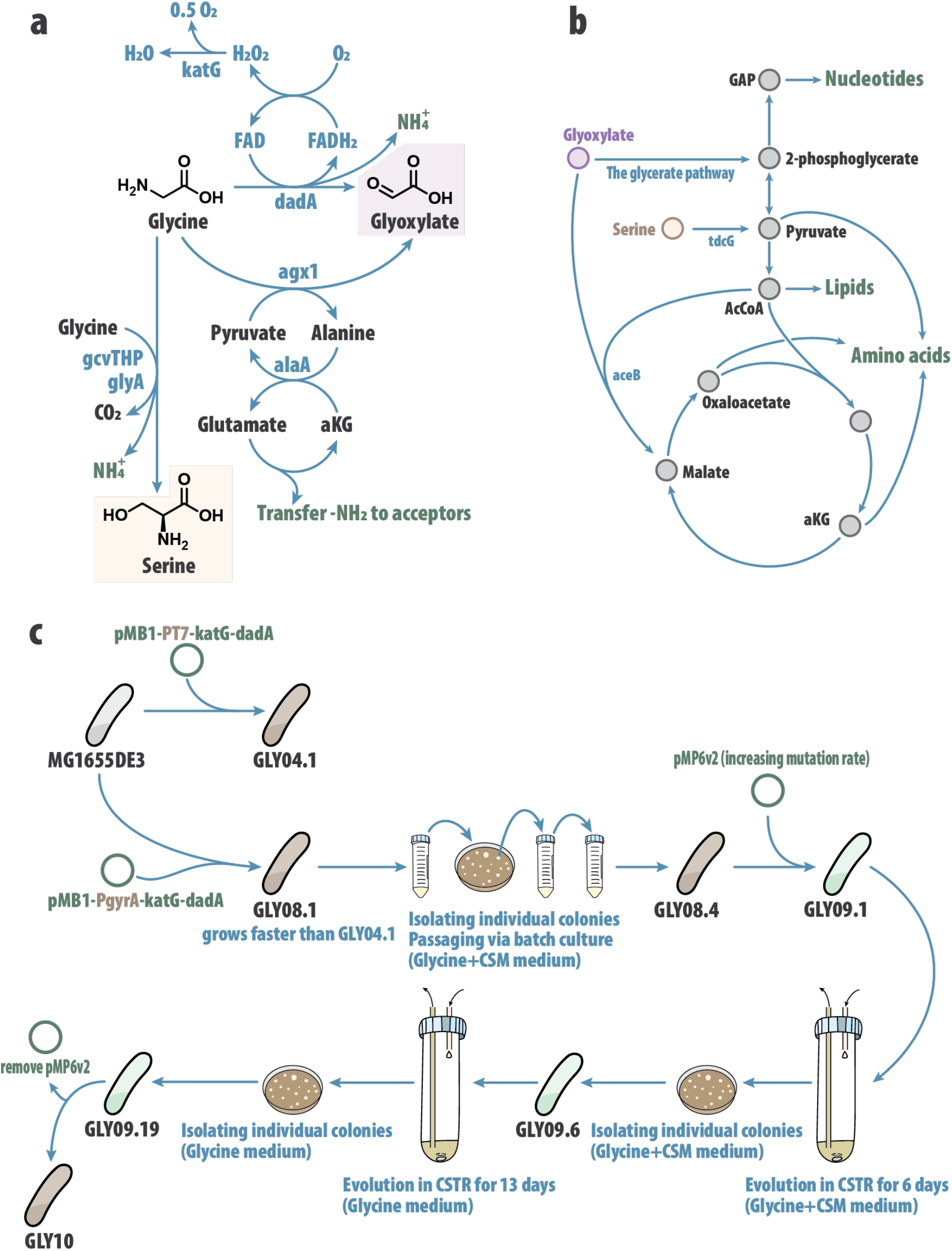
Engineering *E. coli* for glycine utilization. (**a**) Three metabolic pathways of converting glycine into L-serine and glyoxylate in engineered *E. coli*: condensing two glycine into one serine via the glycine cleavage cascade (encoded by *gcvTHP* and *glyA*); transferring the amino group of glycine to pyruvate results in forming glyoxylate (catalyzed by a glycine:alanine transaminase that is encoded by ^Sc^*Agx1*; oxidizing glycine into glyoxylate by a D-amino acid dehydrogenase (encoded by ^Re^*dadA6*). (**b**) Metabolic pathways of converting glyoxylate and L-serine into nucleotides, lipids and amino acids. (**c**) Flowchart for construction and evolution of glycine utilization strains. The final strain **GLY10** was able to grow on glycine as sole carbon source without CSM supplementation. CSM: Complete Supplement Mixture, which is composed of amino acids and nucleobases. GAP: glyceraldehyde 3-phosphate. aKG: alpha-ketoglutarate. *dadA* encodes a D-amino acid dehydrogenase. *katG* encodes a catalase. *agx1* and *alaA* encode transaminases. *gcvTHP* and *glyA* encode enzymes in the glycine cleavage complex.

In this study, we evaluated the three glycine utilization pathways and found that using the D-amino acid oxidase led to a stable *E. coli* strain (**GLY04.1**) which could utilize glycine as the major carbon source with the CSM supplementation (CSM contains a mixture of amino acids and nucleobases). When we changed the T7 promoter driving the over-expression of the D-amino acid oxidase to a constitutive PgyrA promoter, faster cell growth was achieved (**GLY08.1, Figure 1c**). Sequential batch passaging (**GLY08.2 to GLY08.4**) decreased the growth lag phase but the cell still required the CSM supplementation. Next, we introduced a mutagenesis-inducing plasmid (pMP6v2) into **GLY08.4** and evolved the strain (**GLY09.1**) in a continuous stirred tank reactor (CSTR) by gradually increasing the dilution rate. After the first round (6-day) of evolution, we isolated a strain (**GLY09.6**) that could grow on glycine without CSM supplementation. We removed CSM from the minimal media and continued to evolve the strain in CSTR for 13 days to obtain an even faster growing strain (**GLY09.19, Figure 1c**). In the end, we studied the changes in the genome during the batch and continuous evolution to understand how glycine utilization was improved.

## 2. Materials and methods

### 2.1. Chemicals and materials

Glycine, L-serine, L-tyrosine, Tris solution, M9 salts were purchased from Bio Basic Asia Pacific Pte Ltd. Complete Supplement Mixture (CSM) powder was purchased from Sunrise Science Products. Acetonitrile (HPLC-grade), hydrochloric acid (37% purity) and sulfuric acid (98% purity) were purchased from VWR. Sulfuric acid (95% purity) was purchased from J.T. Baker Chemicals. All the other chemicals were purchased from Sigma Aldrich. Commercially available reagents were used as received without purification. All aqueous solutions were prepared using ultrapure water.

### 2.2. Plasmids and strains

The oligonucleotides used in this study were ordered from Integrated DNA Technologies. The full list of plasmids used in this study is provided in **Table 1**, and these plasmids were constructed based on the guanine-thymine (GT) standard-based workflow (Ma et al., 2019). Gene *dadA6* was amplified from *R. eutropha* **H16** genomic DNA. Gene *Agx1* was amplified from *S. cerevisiae* **BY4700** genomic DNA.

**Table 1.** Plasmids and strains used in this study.

| Strains | Plasmids |
| --- | --- |
| GLY01 | pGU01 (pMB1-Spect<sup>R</sup>-P<sub>T7</sub>-gcvTHP-glyA) |
| GLY02 | pGU02 (pMB1-Spect<sup>R</sup>-P<sub>T7</sub>-gcvTHP-glyA-tdcG) |
| GLY03 | pGU03 (pMB1-Spect<sup>R</sup>-P<sub>T7</sub>-Agx1-alaA-tyrB) |
| GLY04 | pGU04 (pMB1-Spect<sup>R</sup>-P<sub>T7</sub>-katG-dadA6) |
| GLY07 | pCT02 (pMB1-Spect<sup>R</sup>-P<sub>T7</sub>-gfp) |
| GLY08 | pGU08 (pMB1-Spect<sup>R</sup>-P<sub>gyrA</sub>-katG-dadA6) |
| GLY09 | pGU08 (pMB1-Spect<sup>R</sup>-P<sub>gyrA</sub>-katG-dadA6) |
|  | pMP6v2 (pSC101-Amp<sup>R</sup>-P<sub>T7</sub>-katG-dadA6) |
| GLY10 | pGU08 (pMB1-Spect<sup>R</sup>-P<sub>gyrA</sub>-katG-dadA6) |
| GLY11 | pGU08 (pMB1-Spect<sup>R</sup>-P<sub>gyrA</sub>-katG-dadA6) |
|  | pGU09 (pSC101-Amp<sup>R</sup>-P<sub>gyrA</sub>-gcvTHP-glyA-tdcG) |
| GLY12 | pGU08 (pMB1-Spect<sup>R</sup>-P<sub>gyrA</sub>-katG-dadA6) |
|  | pGU10 (pSC101-Amp<sup>R</sup>-P<sub>gyrA</sub>-Agx1-alaA-tyrB) |
| GLY13 | pGU08 (pMB1-Spect<sup>R</sup>-P<sub>gyrA</sub>-katG-dadA6) |
|  | pCT03 (pSC101-Amp<sup>R</sup>-P<sub>gyrA</sub>-gfp) |
| GLY14 | pGU08 (pMB1-Spect<sup>R</sup>-P<sub>gyrA</sub>-katG-dadA6) |
|  | pCT01 (pSC101-Amp<sup>R</sup>-P<sub>T7</sub>-gfp) |

*E. coli* MG1655 DE3 ΔrecA ΔendA ΔtyrR ΔpheA DE3 was used as the parent strain. Spect<sup>R</sup>: spectinomycin resistance. Amp<sup>R</sup>: ampicillin resistance. The working concentration of spectinomycin and ampicillin used in cell cultures were 50 µg/mL and 100 µg/mL respectively. gfp: green fluorescence protein, used as a control. In this table, the strains were defined based on the plasmids they harbor. Through evolution, different generations of the strains would be obtained and they are labelled as [Strain name].[Generation number].

The constructed plasmids were transferred via standard electroporation procedure into the parent strain, which is derived from **MG1655DE3**. The transformants were plated on LB agar with appropriate antibiotics (**Table 1**) after incubation at 37 °C overnight unless stated otherwise. The cells harboring plasmids pMP6v2 or pCT01 (**Table 1**) were incubated at 30 °C unless the experiment aimed to purge the plasmids, because they used the temperature sensitive pSC101 replication origin.

After the continuous evolution has ended, pMP6v2 was removed from *E. coli* **GLY09.19** by culturing it in 10 mL of LB with spectinomycin (**Table 1**) at 37 °C overnight in a 50-mL falcon tube. The overnight culture was then diluted by 10^6^-fold with autoclaved ultrapure water and plated on a LB agar plate containing spectinomycin. The plate was incubated at 30 °C overnight. A single colony was randomly picked and cultured in 10 mL of LB with spectinomycin and stored as frozen glycerol stock.

### 2.3. Cell culture and medium

The general workflow for fermentation in this study was as follows: Prior to the fermentation experiment, the *E. coli* strain glycerol stock was used to inoculate 10 mL of LB medium with appropriate antibiotics (**Table 1**) and was incubated in a 50-mL falcon tube at 37 °C/250 rpm overnight. The overnight cell culture was then centrifuged at 3,500 rpm for 5 mins. The overnight grown seed culture was used to inoculate 10 mL of a chemically defined medium (initial optical density at 600 nm [OD_600_]: 0.1). Unless stated otherwise, the *E. coli* culture was then incubated in a 50-mL falcon tube for 72 h (30 °C/250 rpm).

To prepare 10 mL of the chemically defined medium, the following solutions were mixed: 2 mL of the 5x M9 salts solution, 1 mL of glycine stock solution (100 g/L, stock concentration, sterilized by using autoclave), 0.5 mL of L-phenylalanine stock solution (1 g/L, stock concentration, sterilized by using autoclave), 1 µL of IPTG (100 mM, stock concentration, sterilized by using 0.22 µm filter) and 17 µL of the K3 master mix. CSM (final concentration 1 g/L) is supplemented unless stated otherwise. Antibiotic stock solutions were added if needed (more information included in **Table 1**). Ultrapure water was used to top up the volume to 10 mL. The 5x M9 salts solution was prepared by dissolving 56.5 g of M9 salts in 1 L of ultrapure water, and the solution was autoclaved. The composition of the 5x M9 salts solution was 34 g/L Na_2_HPO_4_, 15 g/L KH_2_PO_4_, 2.5 g/L NaCl, 5 g/L NH_4_Cl. K3 master mix was prepared by dissolving 2.5 mL of 0.1 M autoclaved ferric citrate solution, 1 mL of 4.5 g/L autoclaved thiamine solution, 3 mL of 4 mM autoclaved Na_2_MoO_4_ solution, 1 mL of 1 M MgSO_4_ solution, and 1 mL of 1000 X K3 trace elements stock. The 1000 X K3 was prepared by dissolving the following salts in 1 L of water: 5 g of CaCl_2_∙2H_2_O, 1.6 g of MnCl_2_∙4H_2_O, 0.38 g of CuCl_2_∙2H_2_O, 0.5 g of CoCl_2_∙6H_2_O, 0.94 g of ZnCl_2_, 0.03 g of H_3_BO_3_, and 0.4 g of Na_2_EDTA∙2H_2_O. The trace elements stock solution was autoclaved.

### 2.4. Continuous evolution

Continuous evolution experiments were conducted using a customized, low-cost CSTR, which was fabricated based on a round bottom 50 mL polypropylene tube. The tube was placed in a water bath for temperature control. Mixing was achieved using a magnetic stirring bar placed in the tube. The cap of the 50 mL tube (the reactor) was drilled to have openings that were able to tightly fit five silicone tubings, which were used for sampling, air sparging, air exhausting, feeding and harvesting (illustrated in **Figure 4a**). The reactor was autoclaved with a stirring bar inside (the tubings had barbed luer connectors and all of them were capped except the exhaust port). Air was provided using a small 220V AC air pump (EHEIM AIR400). A single-wrapped 0.22 µm syringe filter was used to sterilize the air before it entering the reactor and the flow rate (0.05 L/min) was controlled using a rotameter with control valve. No condenser was used. Two 12V DC peristatic pumps (INTLLAB, built-in tubing size: 1 mm ID and 3 mm OD) were used to control feeding and harvesting. The pumps were controlled using a motor shield (DFRobot Quad DC Motor Driver Shield for Arduino) mounted on an Arduino Mega2560 circuit board, which is connected to a desktop running Windows 10 via a USB cable. The Arduino board was controlled from the desktop using MATLAB. The built-in tubings in the peristatic pumps were sterilized by flowing 200 g/L sodium hydroxide solution, 70 vol% ethanol and autoclaved glycine medium in sequence.

Prior to the continuous evolution experiment, *E. coli* **GLY09.1** strain was used to inoculate 10 mL of LB medium with appropriate antibiotics (**Table 1**) and was incubated in a 50-mL falcon tube at 37 °C/250 rpm overnight. The overnight cell culture was washed and used to inoculate 5 mL of a glycine medium (initial optical density at 600 nm [OD_600_]: 0.1). There were two versions of the medium: with and without CSM as stated in the Results section. The working volume of the bioreactor was fixed at 5 mL. The bioreactor was first operated in batch mode (D = 0 h^−1^) for 24 h. The continuous mode was then turned on, and the dilution rate was adjusted every 24 h. About 1 mL of cell suspension from the bioreactor was withdrawn every 24 h to measure optical density at 600 nm and to quantify glycine concentration. The samples were also examined under a microscope every 24 h to check contamination. At the end of each continuous evolution experiment, the culture was diluted by a 10^6^-fold with autoclaved ultrapure water and was plated on an agar plate which had the same composition as the liquid medium. The agar plate was incubated at 30 °C overnight.

96 colonies were randomly picked from each agar plate and were cultured in the same medium as the CSTR experiment using 96 deep-well culture plate. The culture plate was incubated for 24 h (30 °C/250 rpm). The clone achieving the highest cell density was stored as glycerol stock.

### 2.5. Quantification of cell density and extracellular metabolites

Ten microliter of cell suspension was diluted 20 times with ultrapure water to measure cell density (OD_600_) with a Tecan Infinite M200PRO Microplate reader.

To measure fermentation by-product concentrations, four hundred and fifty microliter of cell suspension was centrifuged at 14,000 rpm for 1 min. The supernatant was then filtered using a 0.22 μm filter (Chemikalie Pte Ltd), and analysed by HPLC (1260 Infinity series HPLC, Agilent), equipped with an Aminex HPX-87H column (300 x 7.8 mm, Bio-Rad). An isocratic flow was used (0.7 mL/min); the mobile phase was 5 mM sulfuric acid (isocratic flow); the column temperature was 50 °C; a refractive index detector (RID) was used to detect fermentation by-products.

Liquid chromatography-mass spectrometry was used to measure glycine concentration without derivatization (Agilent Single-Quadrupole LC/MS). 10 µL of the cell culture was centrifuged at 20,000 g for 2 min. The supernatant was diluted 100 times with M9 minimal medium (without carbon source) containing 0.1 vol% formic acid. The injection volume was 3 µL. Mobile phase A was water containing 0.1 vol% formic acid. Mobile phase B was acetonitrile containing 0.1 vol% formic acid. The column was Discovery HS-F3 column (3.0 x 150 mm, 3 µm, Supelco). The mobile phase gradient program is as follows: 0 min, 100% A; 1 min; 100% A; 2.5 min, 75% A; 4 min, 65% A; 5 min, 5% A; 7.5 min, 5% A; 12.5 min, 100% A; 15 min, 100% A. The column temperature was 40 °C. The ionization method was electrospray, and the mass spectrometry was operated under the positive mode.

### 2.6. Genomic DNA sequencing and RNA sequencing

Genomic DNA extraction and purification were conducted by using the GeneJET Genomic DNA purification kit (ThermoFisher Scientific) in accordance with the manufacturer’s protocol. Equal volume of DNA samples from biological duplicates were mixed. We performed the genomic DNA sequencing using Oxford Nanopore technology in our laboratory in accordance with the manufacturer’s protocols. Genomic DNA from different strains were multiplexed using the Rapid Barcoding Kit (SQK-RBK004) and sequenced using the MinION Flow Cell or Flongle (R9.4.1). The sequencing data were processed by an in-house MATLAB algorithm. The results were validated using the Integrative Genomics Viewer (Robinson et al., 2011).

Total RNA extraction and purification were conducted by using the GeneJET RNA purification kit (ThermoFisher Scientific) in accordance with the manufacturer’s protocol. *E. coli* **GLY08.4** and **GLY10** strains were used to inoculate 10 mL of LB medium with appropriate antibiotics (specified as in **Table 1**) and was incubated in a 50-mL falcon tube at 37 °C/250 rpm overnight. The overnight cell culture was washed and used to inoculate 10 mL of a glycine medium (initial optical density at 600 nm [OD_600_]: 0.1). The glycine medium contained glycine (final concentration: 10 g/L) as the major carbon source. The medium also contained 1 g/L of CSM and 0.05 g/L L-phenylalanine. The experiments were done in biological duplicates. RNA was extracted when OD_600_ reached ~0.5. Equal volume of RNA samples from biological duplicates were mixed. The sequencing provider was NovogeneAIT Genomics. Standard library preparation method for Illumina PE150 was used except no ribosomal RNA depletion was used. The sequencing data were processed by using an in house developed algorithm.

## 3. Results

### 3.1. Establishing a stable glycine-utilizing E. coli strain

The native glycine cleavage system (GcvTHP and GlyA) is the only native pathway in *E. coli* that transforms glycine into L-serine. We confirmed that glycine cannot be utilized by the *E. coli* strains we have in the lab (**MG1655DE3, BL21DE3** and **BW25113**) as a sole carbon source, nor if supplemented with complete supplement mixture (CSM, a mixture of amino acids and nucleobases), although these strains contained the necessary genes of the glycine cleavage system (Okamura-Ikeda et al., 1993). We hypothesized that the transcription level of *gcvTHP* and *glyA* might not be sufficient to sustain cell growth due to *E. coli*’s native complex regulation of these reactions. To test this hypothesis, we over-expressed *gcvTHP* and *glyA* on a plasmid under the control of T7 promoter (pGU01, see **Table 1**). The resulting strain **GLY01.1** failed to grow when 10 g/L of glycine was used with or without 1 g/L of CSM in a chemically defined medium. Since *E. coli* was reported to be unable to utilize L-serine as sole carbon source and high intracellular L-serine concentrations can be detrimental to the cell, a more efficient pathway is required in *E. coli* to assimilate the L-serine into the central metabolism (Mundhada et al., 2016).

In an unpublished work, we found that overexpression of *tdcG* improved *E. coli* growth on L-serine. When we co-expressed *tdcG* (L-serine deaminase) with the glycine cleavage genes (*gcvTHP* and *glyA*), the resulting strain **GLY02.1** required 10 days to grow up to moderate turbidity (Optical Density at 600 nm [OD_600_] = 1.5) when 10 g/L of glycine was used as sole carbon source. With 1 g/L CSM supplementation, the final cell density was more than doubled (OD_600_ = 3.5). Re-passaging the strain to fresh minimal medium yielded similar final turbidity with shorter growth lag phase (48 h) (**Figure S1b**). However, when the cells were streaked on LB agar and four colonies were randomly picked to separately inoculate the same glycine minimal medium with CSM supplementation, all four colonies failed to grow (**Figure S1c**). We hypothesized that a mixed cell population arose, and subpopulations depend on each other to grow on glycine.

We then turned our attention to the other two pathways which can convert glycine into glyoxylate. By over-expressing the glycine:alanine transaminase (encoded by ^Sc^*Agx1*), L-alanine deaminase (encoded by *alaA*) and tyrosine aminotransferase (encoded by *tyrB*) (pGU03), the strain **GLY03.1** managed to grow up to 1.5 OD_600_ units after 96 h when 10 g/L of glycine was used as sole carbon source (**Figure S2a**). **GLY03.1**’s growth can be improved by CSM supplementation, but most isolated colonies cannot grow on glycine, similar to what was observed with **GLY02.1**.

When the D-amino acid dehydrogenase (encoded by ^Re^*dadA6*) and a native catalase (encoded by *katG*) were over-expressed, the strain **GLY04.1** managed to grow up to 1.6 OD_600_ units after 96 h when 10 g/L of glycine was used as carbon source with 1 g/L of CSM supplementation (**Figure 2a**). The catalase was expressed because DadA6’s redox cofactor is FAD, whose recycling reaction was reported to produce hydrogen peroxide (Korshunov & Imlay, 2010).

**Figure 2.**
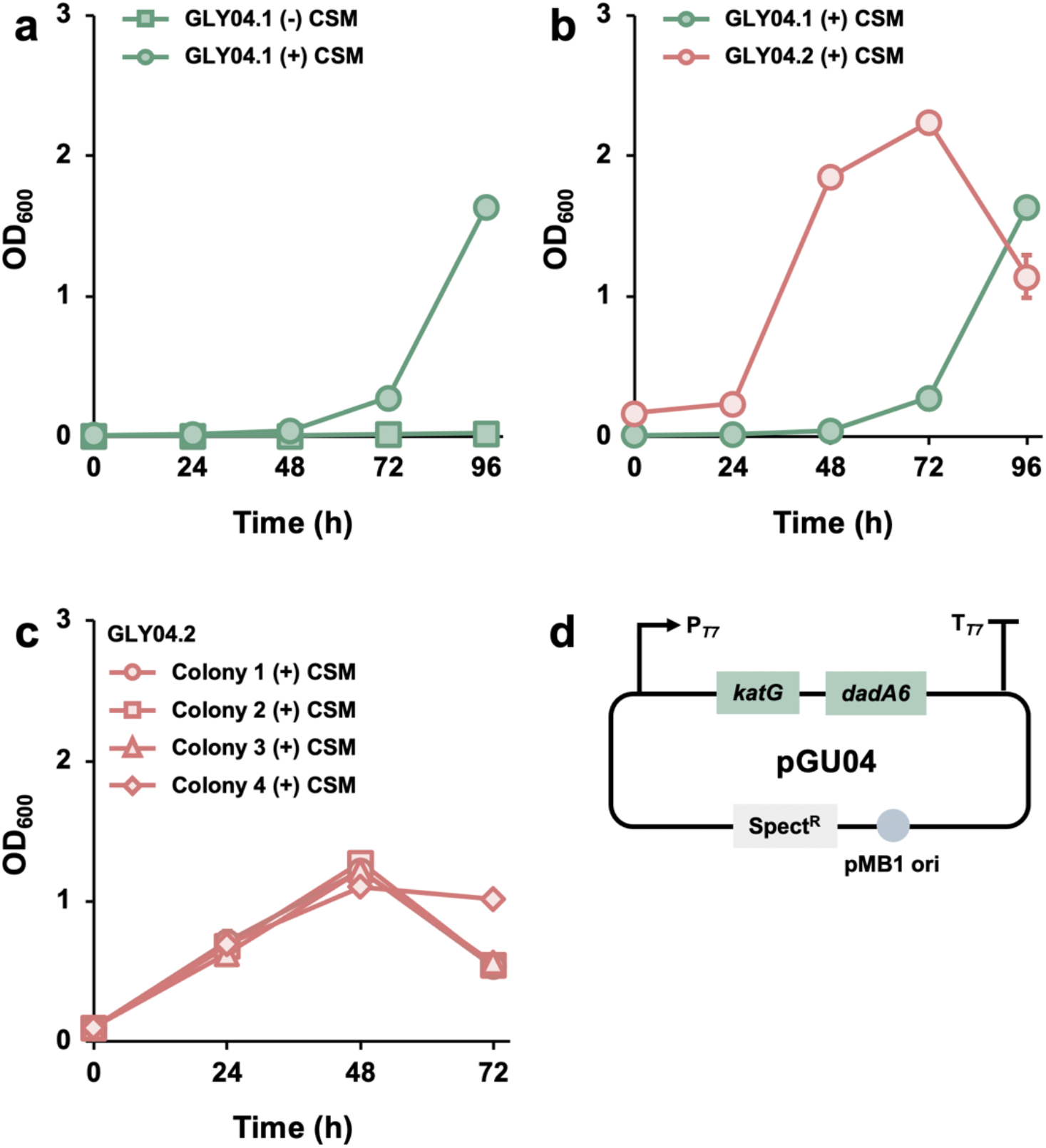
Introducing the D-amino acid dehydrogenase in *E. coli* to utilize glycine. **GLY04** (strain information is in **Table 1**) was cultured in a chemically defined medium containing 10 g/L glycine. The medium was supplemented with 1 g/L CSM and 0.05 g/L L-phenylalanine. (**a**) Cell growth profile of the first-generation strain (**GLY04.1**) with and without CSM. (**b**) Cell growth profile of the first and second-generation strains (**GLY04.1** and **GLY04.2**) with CSM supplementation. (**c**) Cell growth profile of four colonies of the second-generation strain (**GLY04.2**). (**d**) Illustration of the plasmid used by the **GLY04** strains. The size of the error bars (SE, n=2) may be smaller than the symbol sizes.

Without 1 g/L of CSM supplementation, the cells could not grow. Re-passaging **GLY04.1** strain to fresh minimal medium containing 10 g/L of glycine and 1 g/L of CSM yield better growth profile and shorter lag phase. When the cells were streaked on LB agar and all four randomly picked colonies could grow in the liquid medium containing glycine and CSM (**Figure 2c**). Since this pathway allowed us to obtain a stable strain, we next further engineered this pathway.

### 3.2. *Improving glycine utilization in E. coli* through promoter engineering and adaptive laboratory evolution

T7 promoter was used to drive katG-dadA6 in **GLY04.1**. We hypothesized that using T7 promoter may have negative effects on cell growth since it requires addition of IPTG and some cells may not efficiently take up IPTG (Anilionyte et al., 2018). Hence, we replaced the T7 promoter with a constitutive promoter (PgyrA). The resulting strain **GLY08.1** indeed grew at a faster rate than **GLY04.1** with a shorter lag phase (24 h) and an almost doubled maximal cell density (OD_600_ = 3.5, **Figure 3a**). The dependence of CSM supplementation was seen as well, suggesting that amino acid or nucleobases synthesis limited cell growth when glycine served as carbon source. We tested a control where the same basal strain expressed a green fluorescent protein instead of the glycine utilisation gene cassette. The gfp control (**GLY07.1**) could not grow in glycine even when CSM was supplemented, confirming **GLY08.1**’s growth was due to the constructed pathway (**Figure S3a**).

**Figure 3.**
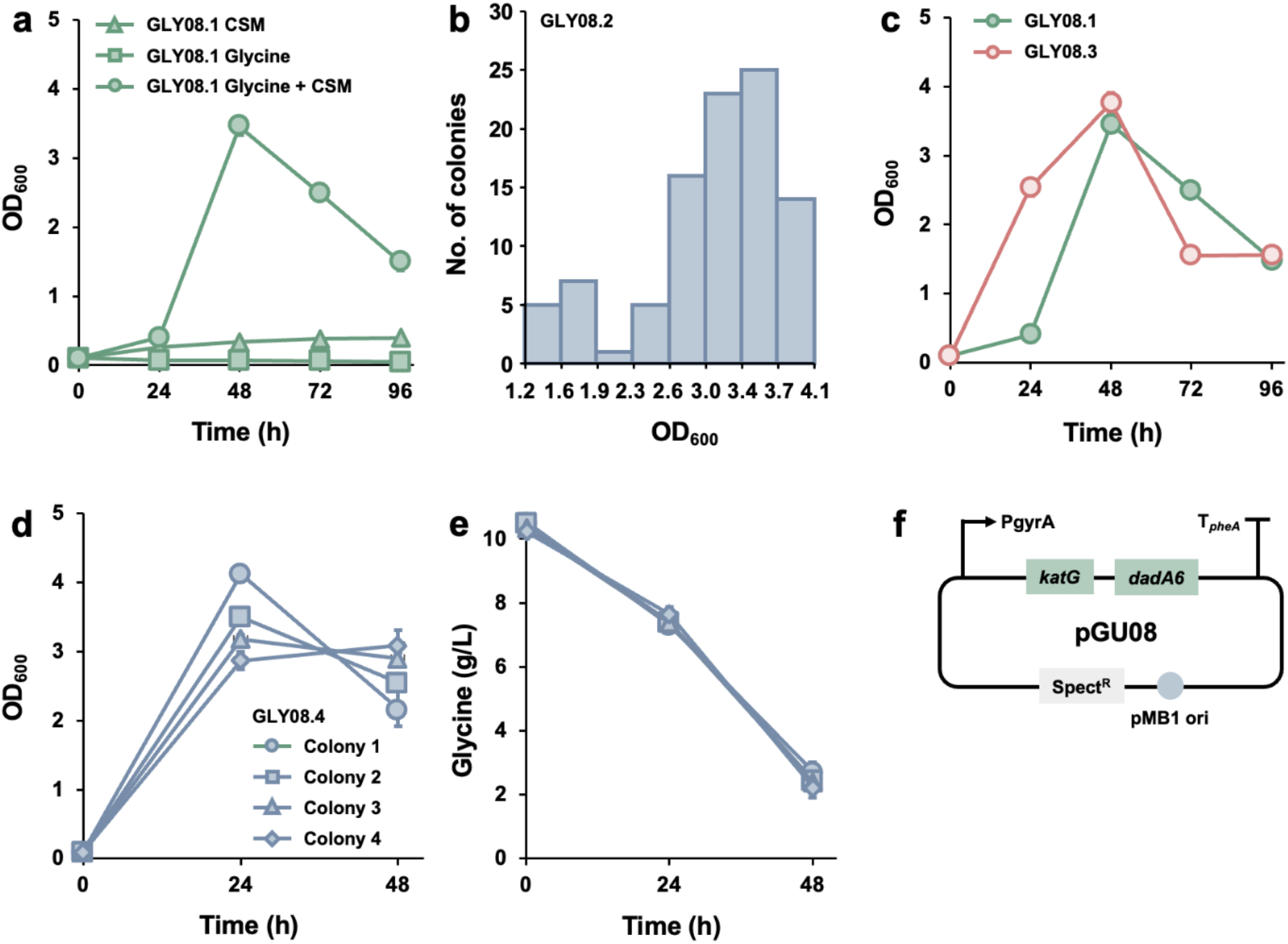
Improving glycine utilisation in *E. coli*. **GLY08** strains were cultured in a chemically defined medium containing 10 g/L glycine, 1 g/L CSM and 0.05 g/L L-phenylalanine. (**a**) Cell growth profile of the first-generation strain (**GLY08.1**). (**b**) Final cell densities of 96 colonies of **GLY08.2** after 48 h culture. (**c**) Comparing cell growth profile of **GLY08.1** and **GLY08.3**. (**d**) Cell growth profile of four randomly picked colonies of **GLY08.4**. (**e**) Glycine concentration (HPLC quantification) were monitored over time. Common fermentation by-products such as acetate and lactate were not detected in the culture media of these fermentations (detection limit: 0.2 g/L). (**f**) Illustration of the plasmid used by the **GLY08** strains. The size of the error bars (SE, n=2) may be smaller than the symbol sizes.

**Figure 4.**
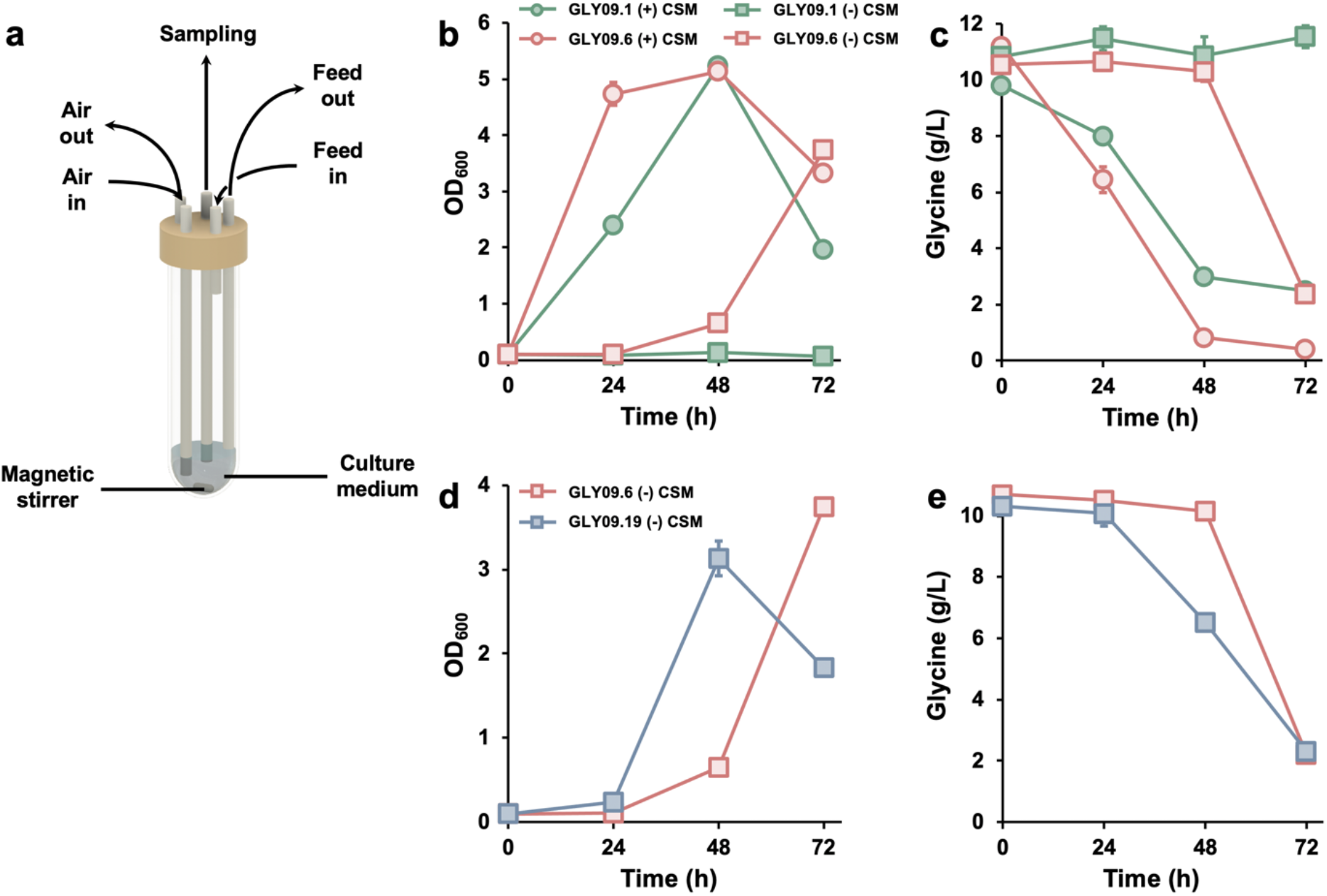
Adaptive laboratory evolution improved glycine utilisation rate and growth rate. **GLY09** (strain information is in **Table 1**) was cultured in a chemically defined medium containing 10 g/L glycine. The medium was supplemented with 1 g/L CSM and 0.05 g/L L-phenylalanine. (**a**) Illustration of the bioreactor vessel to perform constant flow evolution. It was constructed using a 50-mL falcon tube, retrofitted with silicone tubing to provide oxygen and medium transfer. (**b**) Cell growth profile of the first-generation strain (**GLY09.1**) and sixth-generation strain (**GLY09.6**) with and without CSM. (**c**) Glycine concentration (Agilent Single-Quadrupole LC/MS). (**d**) Cell growth profile of the sixth-generation strain (**GLY09.6**) and nineteenth-generation strain (**GLY09.19**) without CSM. (**e**) Glycine concentration (Agilent Single-Quadrupole LC/MS). Common fermentation by-products such as acetate and lactate were not detected in the culture media of these fermentations (detection limit: 0.2 g/L). The size of the error bars (SE, n=2) may be smaller than the symbol sizes.

To determine if **GLY08.1**’s cell population was mostly homogenous, we plated the strain on LB agar. 96 colonies were grown using deep well plate. After 48 h incubation, the second-generation strains exhibited different final cell densities (**Figure 3b**). We then chose the strain which exhibited the highest final cell density (OD_600_ = 4.1, **GLY08.2**) for further passaging into fresh medium with glycine and CSM. The third-generation strain (**GLY08.3**) exhibited better growth profile than the first-generation as the lag phase was shortened (**Figure 3c**). The maximal cell density achieved, and the glycine consumption profile were similar to those of **GLY08.1** (OD_600_ = 3.8, 8 g/L of glycine was consumed, **Figure 3d-e**).

To further improve the growth rate of **GLY08.4**, we introduced a mutator plasmid for speeding up mutagenesis rate in *E. coli*. The plasmid (pMP6v2) was derived from pMP6 (Badran & Liu, 2015) by changing the promoter from pBAD to pT7 and replacing the CloDF13 replication origin with a temperature-sensitive version of pSC101. The pBAD promoter is induced by arabinose, which could serve as a carbon source for *E. coli* and thus may interfere with glycine utilization. The new replication origin could allow us to easily remove the plasmid when needed. After introducing pMP6v2 into **GLY08.4**, the resulting strain **GLY09.1** was evolved in a continuous stirred tank reactor (CSTR) by using leaky pT7 expression (adding IPTG was found to negatively affect cell growth, **Figure S4a**). The CSTR was built with commonly available lab consumables and was designed to have a small reaction volume (5 mL) to conserve medium usage (**Figure 4a**). By increasing the dilution rate incrementally, the mutants exhibiting a specific growth rate lower than dilution rate would be washed out, thus selecting for the mutants exhibiting a higher specific growth rate. After evolving **GLY09.1** for six days, we isolated a strain (**GLY09.6**) which grew faster than **GLY09.1** (**Figures 4b** and **S5b**). The specific growth rate increased from 0.11 h^−1^ (**GLY09.1**) to 0.17 h^−1^ (**GLY09.6**). The overall consumption of glycine at 72 h was also improved from 75% (**GLY09.1**) to 99% (**GLY09.6**). This mutant strain is also able to grow in glycine without CSM supplementation (**Figure 4b-c**). We then further evolved **GLY09.6** in a CSTR for 13 days, in which CSM was omitted from the minimal medium feed. We isolated a strain (**GLY09.19**) which had a much shorter lag phase than **GLY09.6** (**Figure 4d** and **S6b**). The specific growth rate of this mutant (0.07 h^−1^) was slightly higher than that of **GLY09.6** (0.05 h^−1^).

### 3.3. Characterization of the evolved strains

To obtain a stable strain (**GLY10**), we removed pMP6v2 from **GLY09.19** by growing the cells at 37 °C (the cells were grown at 30 °C during the evolution). **GLY10** was able to consume 8 g/L of glycine within 48 h without the CSM supplementation (**Figure 5**).

**Figure 5.**
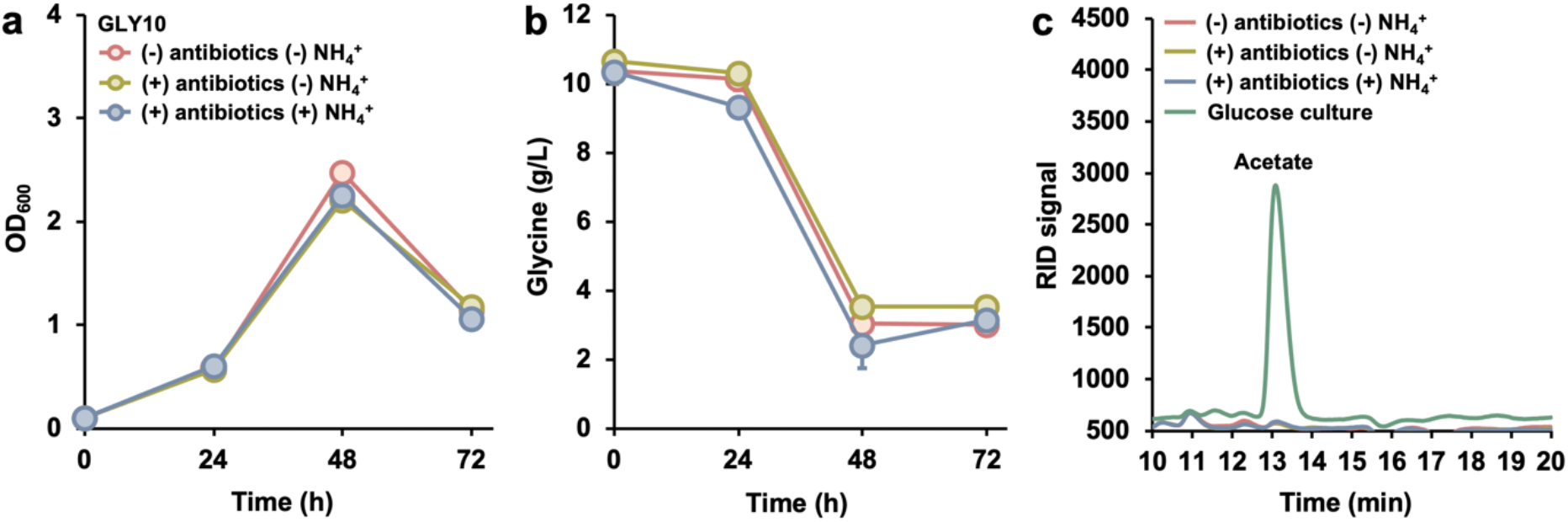
Effects of removing ammonium and antibiotics on cell growth and glycine consumption of GLY10. The strain was cultured in a chemically defined medium containing 10 g/L glycine and 0.05 g/L L-phenylalanine. Cell growth profile (**a**) and glycine profile (**b**) of **GLY10** with and without spectinomycin/ammonium chloride. (**c**) Chromatograms (HPLC RID) of glycine and glucose culture at 72 h. **GLY10** was grown in the glucose medium (the same medium recipe as the glycine medium except 10 g/L glycine was replaced with 10 g/L glucose) as a control to demonstrate that **GLY10** did not produce fermentation by-products. The size of the error bars (SE, n=2) may be smaller than the symbol sizes.

In all the above-described experiments, the culture media contained 1 g/L ammonium chloride as part of the standard M9 medium recipe, and spectinomycin was used to maintain the pGU08 plasmid. Since the amino group of glycine was released as ammonium ion during glycine’s assimilation, we hypothesized that the ammonium salt in the medium can be omitted. 10 g/L glycine can release 2.4 g/L ammonium ion, which is equivalent to that provided by 7.1 g/L ammonium chloride and much higher than that included in the M9 salts. We also hypothesized that spectinomycin was not needed by the strain, because any cell losing the plasmid should not grow on glycine. We omitted ammonium chloride and spectinomycin in the medium formulation, and indeed found that the cell growth and glycine consumption were not affected (**Figure 5a-b**). Glycine was used as the sole carbon and nitrogen source to support the growth of **GLY10**. Acetate and other *E. coli* fermentation by-products were not detected through the entire process (detection limit: 0.2 g/L, **Figure 5c**).

To understand the mechanisms underlying the adaptive evolution, we sequenced the genome of **GLY08.1** (the starting strain), **GLY08.4** (after batch passaging and before introducing pMP6v2 and starting the CSTR evolution) and the final strain (**GLY10**). While only one new nucleotide polymorphism (SNP) was found in **GLY08.4** compared with **GLY08.1**, 40 new SNPs were detected in **GLY10** using **GLY08.4** as the reference, and 92% of the SNPs were found in the coding region. This suggests that pMP6v2 was highly efficient in increasing the genome mutagenesis rate during the continuous evolution (**Table S1**).

Among the 40 SNPs, the one causing the F158L mutation in glutamate synthase GltB should be highlighted, given the importance of the enzyme in nitrogen metabolism and ammonia assimilation. It may have played a key role in enabling **GLY10** to grow without CSM. The second SNP that caught our attention was the M26I mutation in *atpG*, which is a critical component in the ATP synthase F1 complex (Tang & Capaldi, 1996). This mutation could affect the efficiency of the cellular energy harvesting from the proton translocation across the inner membranes. The last two we highlight are related to mRNA synthesis (S156P in NusG, transcription termination factor) and degradation (E132K in RhlB, ATP-dependent RNA helicase) (Burns & Richardson, 1995; Coburn et al., 1999; Py et al., 1996). Mutations in such enzymes may affect transcription of a large number of genes.

To further investigate the mechanism, we quantified the transcriptome of **GLY08.4** and **GLY10** during their early exponential growth phase in the glycine+CSM medium (10 g/L glycine and 1 g/L CSM) (**Table S2**). The results suggest that the cell fine-tuned the gene expressions around the glyoxylate node during the evolution to reduce the accumulation of glyoxylate, which is a toxic aldehyde intermediate. DadA6 produces glyoxylate from glycine. Its transcription level in **GLY10** was only 17% of that in **GLY08.4**. The glycerate pathway is an essential pathway for assimilating glyoxylate into the central metabolism. All the genes involved in the glycerate pathway were substantially upregulated when **GLY08.4** evolved into **GLY10** (fold change: *gcl*: 2.9, *glxR*: 3.5, *glxK*: 3, *garK*: 5.9, **Figure 6**). Glyoxylate could also be assimilated into the TCA cycle when borrowing acetyl-CoA or succinate. The genes controlling these two secondary glyoxylate assimilation pathways were upregulated too (fold change: *aceB*: 3.6, *glcB*: 4, *aceA*: 7.1, *aceK*: 5.6). The house-keeping aldehyde dehydrogenase (AldA) was upregulated by a factor of 6.2 and two aldehyde dehydrogenase genes were activated (the fold change: *astD*: 107, *aldB*: 102). These dehydrogenases could have played roles in detoxification of glyoxylate and other aldehyde intermediates.

**Figure 6.**
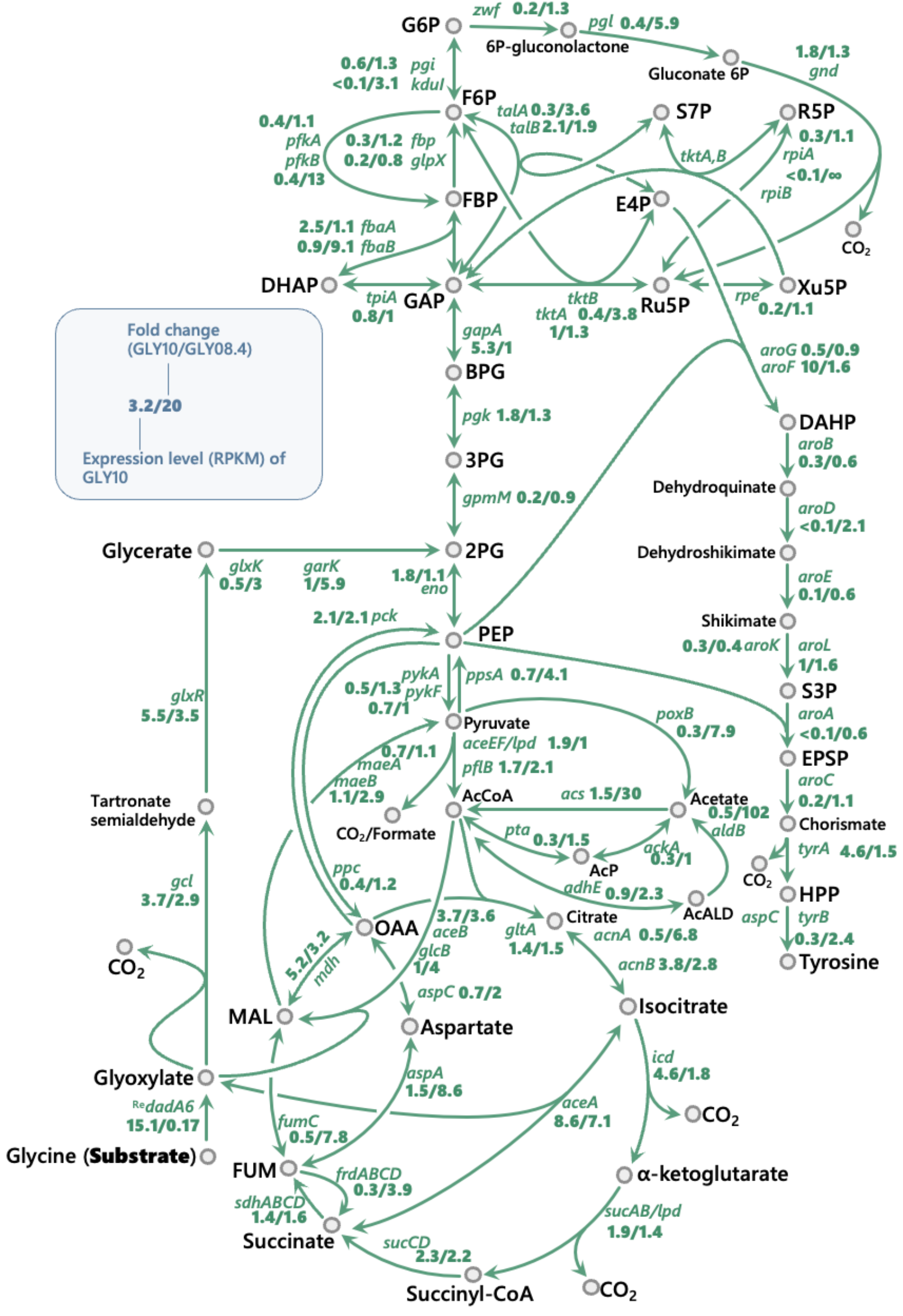
Relative transcriptome analysis of GLY08.4 and GLY10 strain when growing in a glycine medium. The strain was cultured in a chemically defined medium containing 10 g/L glycine, 1 g/L CSM, and 0.05 g/L phenylalanine. RNA was extracted when OD600 reached ~0.5. Equal quantity of RNA from two biological duplicates were mixed and sent to service provider for sequencing. The expression level was provided in standard RPKM values.

During the evolution, there were obvious upregulations in pentose phosphate pathway (the fold change: *pgl*: 5.9; *talA*: 3.6; *tktA*: 3.8), through which the cell may increase the NADPH regeneration rate or increase the flux towards nucleotide synthesis and/or aromatic amino acid production. The cell has also activated/enhanced two pathways to transform pyruvate into acetyl-CoA: the fold change of the formate dehydrogenase was 2.1 and the two genes in the acetate bypass were upregulated (the fold change: *poxB*: 7.9; *acs*: 30). Increasing the availability of acetyl-CoA could allow the cells to the acetyl-CoA-dependent pathway to assimilate glyoxylate into the TCA cycle (through malate synthases AceB and GlcB).

## 4. Discussion

We have successfully engineered an *E. coli* strain to utilize glycine as sole carbon and nitrogen source. It was based on the glycine dehydrogenase pathway. Using the native glycine cleavage system did not yield stable strains, possibly due to the tight regulations of the enzymes by the host. The glyoxylate-alanine transaminase pathway also failed to work reliably under the conditions we tested. Our hypothesis is that the transamination cascade could not process the large amount of amino group from glycine. Such overwhelming influx of amino group could deplete the acceptor, pyruvate, which plays central role in the host metabolism. The successfully implemented dehydrogenase pathway releases the amino group as ammonium ions to the media so it would not be subject to this constraint. We have briefly explored using the dehydrogenase pathway together with the glycine cleavage system or the transaminase system, but neither yielded significant improvements (**Figures S7a-b**).

The CSTR was useful in selecting mutants that have higher specific growth rate. Its operation is fully automated, so it is more friendly to the user compared with tedious batch passaging. The continuous operation is also more effective because the cells are always in their growing phase, without the need of adapting to fresh medium. Most commercially available reactors are too large for continuous evolution application as it would require large amounts of feed. They are also expensive instruments to many labs. In this study, our CSTR was built using common laboratory consumables, low-cost peristatic pumps and Arduino circuit boards. The system had a small reaction volume (5 mL) and was controlled using MATLAB with a graphical user interface.

The mutagenesis-inducing plasmid was important in introducing mutations to *E. coli*’s genome to speed up the evolution process. Unlike traditional methods which employs UV irradiation or toxic mutagenic chemicals, the mutator plasmid is safe to handle as it will not pose any hazard to human health. Moreover, we found that the leaky expression of pMP6v2 was sufficient to express mutagenic elements without compromising cell growth. We believe the combination of the mutagenesis-inducing plasmid and CSTR can be used in other studies such as improving formate utilisation in *E. coli*. The current best engineered strain (**GLY10**) may be further engineered for this purpose since glycine is a key intermediate in the formate utilisation pathway.

## Author contributions

**Vincent Fung:** Conceptualization, Methodology, Investigation, Formal analysis, Writing – original draft. **Daniel Tan:** Investigation, Writing – original draft. **Kang Zhou:** Conceptualization, Methodology, Supervision, Writing – review & editing, Funding acquisition.

## Conflict of interest statement

The authors declare no competing interest.

## Acknowledgement

This work is financially supported by Ministry of Education (MOE) Singapore through a Tier-2 research grant (identifier: R-279-000-594-112). V.F. was financially supported by PhD scholarships from National University of Singapore (NUS) and MOE Singapore. We acknowledge useful guidance from Dr. Liming Yang on using analytical instruments in the Department of Chemical & Biomolecular Engineering, NUS.

## Data availability

All data needed to evaluate the conclusions in the paper are present in the paper and/or the Supplementary information.

## Notes

### Competing Interest Statement

The authors have declared no competing interest.

## References

Anilionyte, O., Liang, H., Ma, X., Yang, L., & Zhou, K. (2018). Short, auto-inducible promoters for well-controlled protein expression in Escherichia coli. Applied Microbiology and Biotechnology, 102(16), 7007–7015. https://doi.org/10.1007/s00253-018-9141-z

Badran, A. H., & Liu, D. R. (2015, 2015/10/07). Development of potent in vivo mutagenesis plasmids with broad mutational spectra. Nature Communications, 6(1), 8425. https://doi.org/10.1038/ncomms9425

Burman, J. D., Harris, R. L., Hauton, K. A., Lawson, D. M., & Sawers, R. G. (2004). The iron–sulfur cluster in the l-serine dehydratase TdcG from Escherichia coli is required for enzyme activity. FEBS letters, 576(3), 442–444. https://doi.org/10.1016/j.febslet.2004.09.058

Burns, C. M., & Richardson, J. P. (1995). NusG is required to overcome a kinetic limitation to Rho function at an intragenic terminator. Proceedings of the National Academy of Sciences, 92(11), 4738–4742. https://doi.org/10.1073/pnas.92.11.4738

Claassens, N. J., Bordanaba-Florit, G., Cotton, C. A. R., De Maria, A., Finger-Bou, M., Friedeheim, L., Giner-Laguarda, N., Munar-Palmer, M., Newell, W., Scarinci, G., Verbunt, J., de Vries, S. T., Yilmaz, S., & Bar-Even, A. (2020, 2020/11/01/). Replacing the Calvin cycle with the reductive glycine pathway in Cupriavidus necator. Metabolic Engineering, 62, 30–41. https://doi.org/https://doi.org/10.1016/j.ymben.2020.08.004

Coburn, G. A., Miao, X., Briant, D. J., & Mackie, G. A. (1999). Reconstitution of a minimal RNA degradosome demonstrates functional coordination between a 3’ exonuclease and a DEAD-box RNA helicase. Genes & Development, 13(19), 2594–2603. https://doi.org/10.1101/gad.13.19.2594

Korshunov, S., & Imlay, J. A. (2010). Two sources of endogenous hydrogen peroxide inEscherichia coli. Molecular Microbiology, 75(6), 1389–1401. https://doi.org/10.1111/j.1365-2958.2010.07059.x

Ma, X., Liang, H., Cui, X., Liu, Y., Lu, H., Ning, W., Poon, N. Y., Ho, B., & Zhou, K. (2019, 2019/07/23). A standard for near-scarless plasmid construction using reusable DNA parts. Nature Communications, 10(1), 3294. https://doi.org/10.1038/s41467-019-11263-0

Maia, L. B., Moura, I., & Moura, J. J. G. (2021). Carbon Dioxide Utilisation—The Formate Route. In Enzymes for Solving Humankind’s Problems (pp. 29–81). https://doi.org/10.1007/978-3-030-58315-6_2

Mundhada, H., Schneider, K., Christensen, H. B., & Nielsen, A. T. (2016, Apr). Engineering of high yield production of L-serine in Escherichia coli. Biotechnol Bioeng, 113(4), 807–816. https://doi.org/10.1002/bit.25844

Neuendorf, C. S., Vignolle, G. A., Derntl, C., Tomin, T., Novak, K., Mach, R. L., Birner-Grünberger, R., & Pflügl, S. (2021). A quantitative metabolic analysis reveals Acetobacterium woodii as a flexible and robust host for formate-based bioproduction. Metabolic Engineering, 68, 68–85. https://doi.org/10.1016/j.ymben.2021.09.004

Newman, E. B., & Walker, C. (1982). L-serine degradation in Escherichia coli K-12: a combination of L-serine, glycine, and leucine used as a source of carbon. Journal of Bacteriology, 151(2), 777–782. https://doi.org/10.1128/jb.151.2.777-782.1982

Okamura-Ikeda, K., Ohmura, Y., Fujiwara, K., & Motokawa, Y. (1993). Cloning and nucleotide sequence of the gcv operon encoding the Escherichia coli glycine-cleavage system. European journal of biochemistry, 216(2), 539–548. https://doi.org/10.1111/j.1432-1033.1993.tb18172.x

Py, B., Higgins, C. F., Krisch, H. M., & Carpousis, A. J. (1996). A DEAD-box RNA helicase in the Escherichia coli RNA degradosome. Nature, 381(6578), 169–172. https://doi.org/10.1038/381169a0

Robinson, J. T., Thorvaldsdóttir, H., Winckler, W., Guttman, M., Lander, E. S., Getz, G., & Mesirov, J. P. (2011). Integrative genomics viewer. Nature Biotechnology, 29(1), 24–26. https://doi.org/10.1038/nbt.1754

Su, H., & Newman, E. B. (1991). A novel L-serine deaminase activity in Escherichia coli K-12. Journal of Bacteriology, 173(8), 2473–2480. https://doi.org/10.1128/jb.173.8.2473-2480.1991

Su, H. S., Lang, B. F., & Newman, E. B. (1989). L-serine degradation in Escherichia coli K-12: cloning and sequencing of the sdaA gene. Journal of Bacteriology, 171(9), 5095–5102. https://doi.org/10.1128/jb.171.9.5095-5102.1989

Takada, Y., & Noguchi, T. (1985, Oct 1). Characteristics of alanine: glyoxylate aminotransferase from Saccharomyces cerevisiae, a regulatory enzyme in the glyoxylate pathway of glycine and serine biosynthesis from tricarboxylic acid-cycle intermediates. Biochem J, 231(1), 157–163. https://doi.org/10.1042/bj2310157

Tang, C., & Capaldi, R. A. (1996). Characterization of the Interface between γ and ε Subunits of Escherichia coli F1-ATPase. Journal of Biological Chemistry, 271(6), 3018–3024. https://doi.org/10.1074/jbc.271.6.3018

Umbarger, H. E., & Brown, B. (1957). Threonine deamination in Escherichia coli. II. Evidence for two L-threonine deaminases. Journal of Bacteriology, 73(1), 105–112. https://doi.org/10.1128/jb.73.1.105-112.1957

Van Peteghem, L., Sakarika, M., Matassa, S., Pikaar, I., Ganigué, R., & Rabaey, K. (2022). Towards new carbon–neutral food systems: Combining carbon capture and utilization with microbial protein production. Bioresource Technology, 349. https://doi.org/10.1016/j.biortech.2022.126853

Yu, H., & Liao, J. C. (2018, 2018/09/28). A modified serine cycle in Escherichia coli coverts methanol and CO2 to two-carbon compounds. Nature Communications, 9(1), 3992. https://doi.org/10.1038/s41467-018-06496-4

