## Supplementary material for "Improving glycine utilization in *Escherichia coli*": Glycine SI.pdf

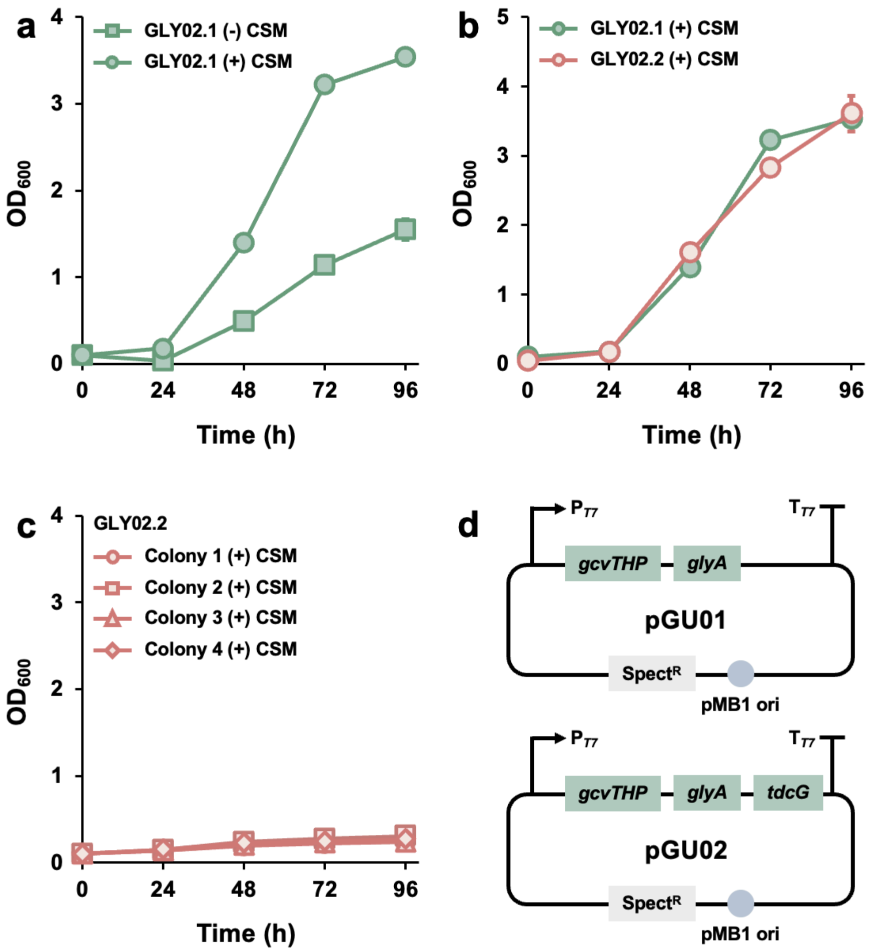


**Figure S1 Introducing the glycine cleavage system in *E. coli* to utilize glycine. GLY02** (strain information is in **Table 1**) was cultured in a chemically defined medium containing 10 g/L glycine. The media was supplemented with 1 g/L CSM and 0.05 g/L L-phenylalanine. (**a**) Cell growth profile of the first-generation strain (**GLY02.1**) with and without CSM**.** (**b**) Cell growth profile of the first and second-generation strain (**GLY02.1** and **GLY02.2**) with CSM supplementation. (**c**) Cell growth profile of four random colonies of the second-generation strain (**GLY02.2**) with CSM supplementation. (**d**) Illustration of the plasmids used by the **GLY02** strains. The size of the error bars (SE, n=2) may be smaller than the symbol sizes.

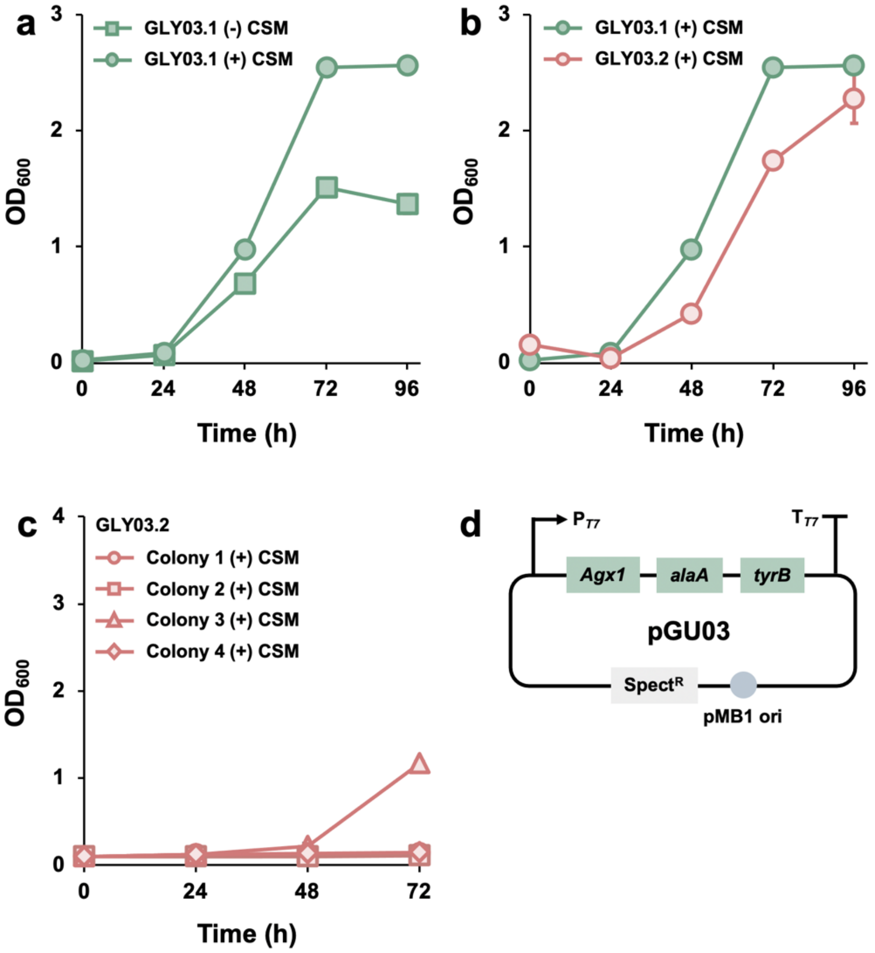


**Figure S2 Introducing the glycine:alanine transaminase system in *E. coli* to utilize glycine. GLY03** (strain information is in **Table 1**) was cultured in a chemically defined medium containing 10 g/L glycine. The media was supplemented with 1 g/L CSM and 0.05 g/L L-phenylalanine. (**a**) Cell growth profile of the first-generation strain (**GLY03.1**) with and without CSM. (**b**) Cell growth profile of the first and second-generation strain (**GLY03.1** and **GLY03.2**) with CSM supplementation. (**c**) Cell growth profile of four random colonies of the second-generation strain (**GLY03.2**) with CSM supplementation. (**d**) Illustration of the plasmid used by the **GLY03** strains. The size of the error bars (SE, n=2) may be smaller than the symbol sizes.

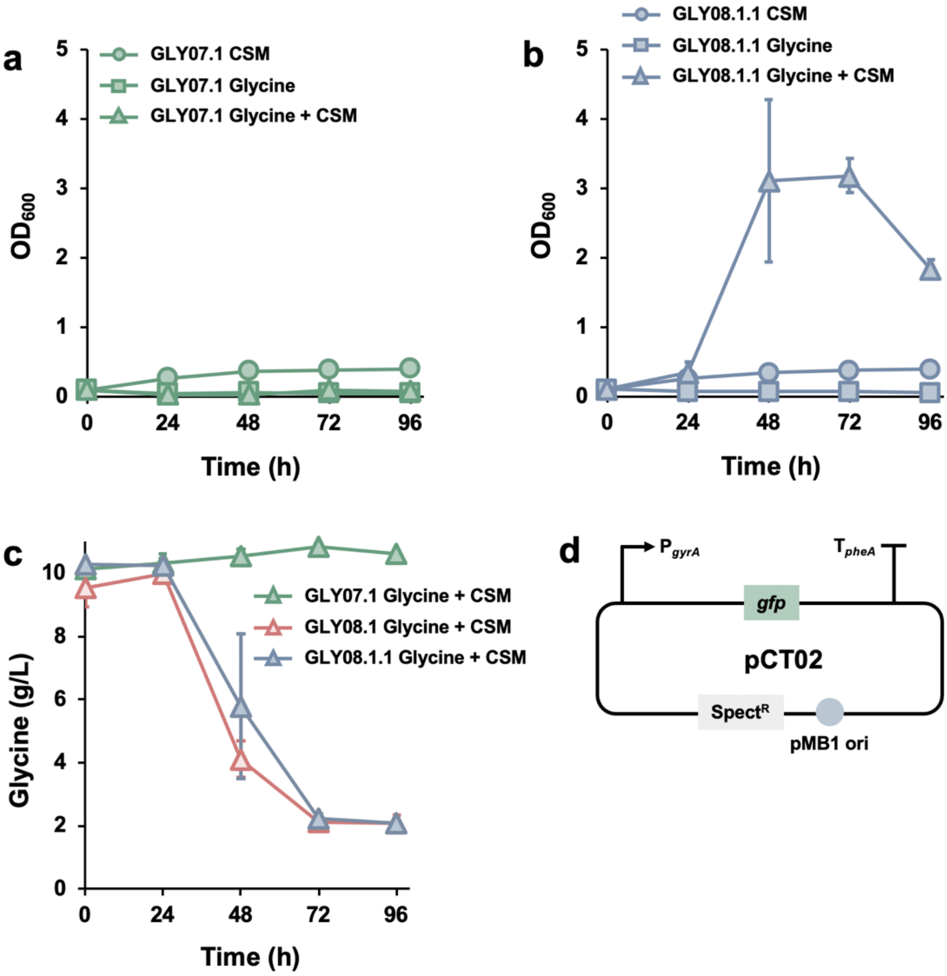


**Figure S3 Investigating if the improved growth rate in glycine culture was due to plasmid mutations. GLY08** (strain information is in **Table 1**) was cultured in a chemically defined medium containing 10 g/L glycine. The media was supplemented with 1 g/L CSM and 0.05 g/L L-phenylalanine. (**a**) Cell growth profile of control strain (**GLY07.1**). (**b**) Cell growth profile of **GLY08.1.1** which contains the plasmid extracted from the fourth-generation strain (**GLY08.4**). (**c**) Glycine concentration (HPLC quantification) were monitored over time. Common fermentation by-products such as acetate and lactate were not detected in the culture media of these fermentations (detection limit: 0.2 g/L). (**d**) Illustration of the plasmid used by the **GLY07** strain. The size of the error bars (SE, n=2) may be smaller than the symbol sizes.

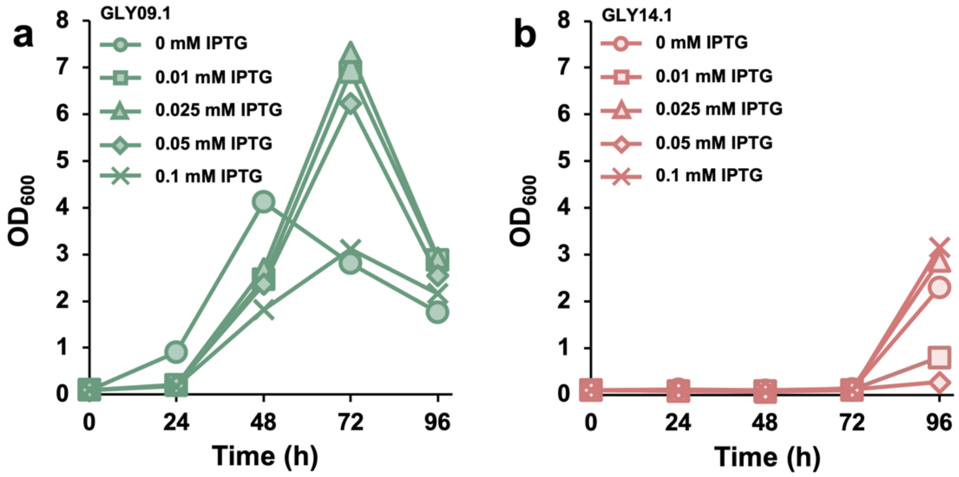


**Figure S4 Introducing the mutagenesis plasmid to improve *E. coli*'s growth rate in glycine culture.** **GLY09** and **GLY14** (strain information is in **Table 1**) was cultured in a chemically defined medium containing 10 g/L glycine. The media was supplemented with 1 g/L CSM and 0.05 g/L L-phenylalanine. (**a**) Cell growth profile of **GLY09.1** strain with CSM supplementation. (**b**) Cell growth profile of **GLY14.1** strain with CSM supplementation.

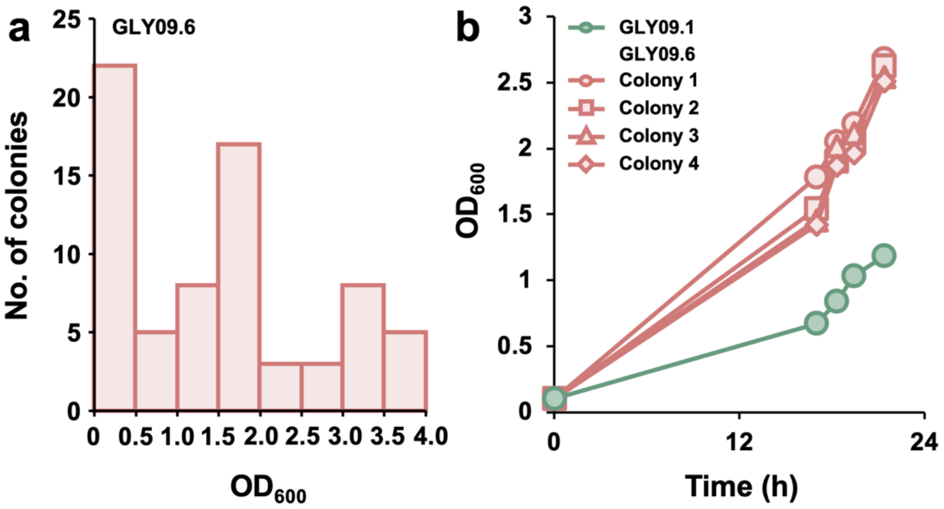


**Figure S5 Adaptive laboratory evolution using the continuous stirred tank reactor (CSTR).** **GLY09.1** was cultured in a chemically defined medium containing 10 g/L glycine. The media was supplemented with 1 g/L CSM and 0.05 g/L L-phenylalanine. (**a**) Final cell densities of 71 colonies of **GLY09.6** after 24 h culture. (**b**) Comparing the growth profile between four isolated strains after 6 days of CSTR evolution (**GLY09.6**) and a strain before CSTR evolution (**GLY09.1**) in the presence of CSM.

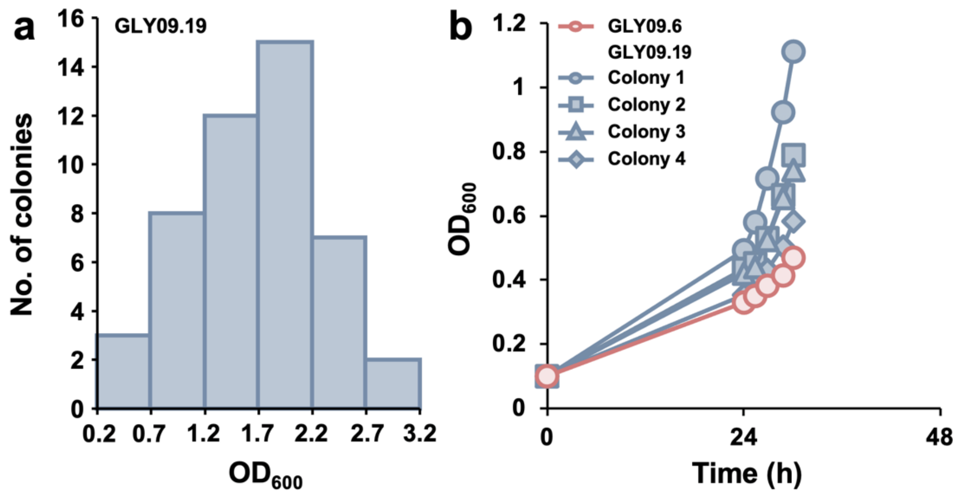


**Figure S6 Adaptive laboratory evolution using the continuous stirred tank reactor (CSTR). GLY09.6** was cultured in a chemically defined medium containing 10 g/L glycine. The media was supplemented with 0.05 g/L L-phenylalanine. (**a**) Final cell densities of 47 colonies of **GLY09.19** after 48 h culture without CSM supplementation. (**b**) Comparing the growth profile between four isolated strains after 13 days of CSTR evolution (**GLY09.19**) and a strain before CSTR evolution (**GLY09.6**) in the absence of CSM.

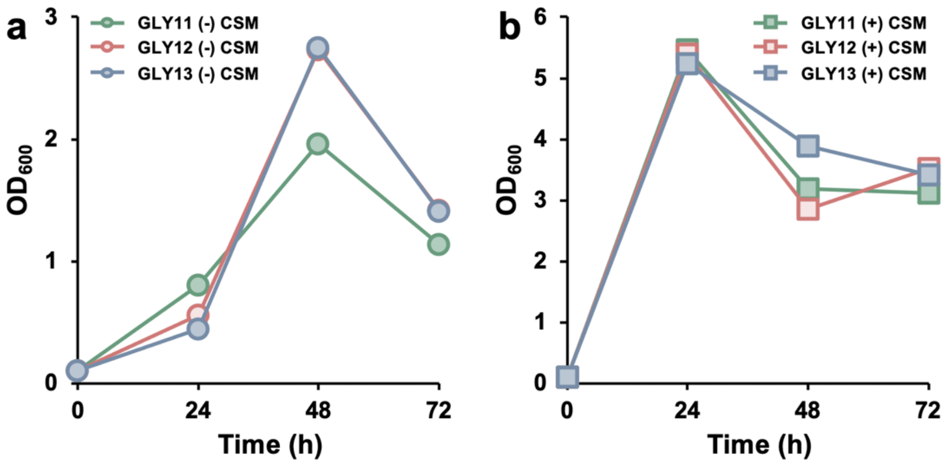


**Figure S7 Introducing multiple glycine utilisation pathways to improve *E. coli*'s growth rate in glycine culture. GLY11, GLY12** and **GLY13** (strain information is in **Table 1**) was cultured in a chemically defined medium containing 10 g/L glycine. The media was supplemented with 1 g/L CSM and 0.05 g/L L-phenylalanine. (**a**) Cell growth profile of **GLY11**, **GLY12** and **GLY13** strain without CSM supplementation. (**b**) Cell growth profile of **GLY11**, **GLY12** and **GLY13** strain with CSM supplementation. The size of the error bars (SE, n=2) may be smaller than the symbol sizes.
